# Near-complete, haplotype-resolved genome assembly of common buckwheat (*Fagopyrum esculentum* Moench)

**DOI:** 10.64898/2026.03.30.715208

**Authors:** Fabian Hess, Yutang Chen, María-Estefanía López, Axelle Colliquet, Ingrid Stoffel-Studer, Victor Mac, Stefan Grob, Roland Kölliker, Bruno Studer

**Author notes:** Corresponding author: Bruno Studer. These authors contributed equally (Fabian Hess, Yutang Chen).

## Abstract

Common buckwheat (*Fagopyrum esculentum* Moench) is a globally cultivated pseudocereal with a high nutritional quality and economic value. Due to its self-incompatibility, common buckwheat exhibits a high level of heterozygosity, making genome assembly challenging. Consequently, reference-level haplotype-resolved assemblies of common buckwheat are scarce, hindering research and genomics-assisted breeding. Here, we present a near-complete, chromosome-level, haplotype-resolved assembly of a common buckwheat F_1_ genotype (named Tuka), generated using a trio-binning approach that integrated parental Illumina short-read data with PacBio HiFi and Hi-C data from Tuka. The Tuka assembly comprises two haplomes, Tuka_h1 and Tuka_h2, both showing high contiguity (contig N50 of 76.68 Mb and 84.57 Mb, respectively), high completeness (assembly sizes of 1.28 Gb and 1.23 Gb with BUSCO scores of 96.9% and 96.8%, respectively), high base-level accuracy (QV of 59.08 and 63.03, respectively), and few gaps (35 and 30, respectively). This near-complete assembly of Tuka serves as a valuable genomic resource for common buckwheat, enabling advanced genomic analyses and accelerating research and breeding using state-of-the-art genomic tools.

## Background & Summary

Common buckwheat (*Fagopyrum esculentum* Moench) is an underutilized crop with a great potential to diversify agroecosystems and contribute to more resilient and productive food systems^1-3^. However, as an underutilized crop, its agronomic performance generally lags behind that of major crops, indicating the need for concerted breeding efforts to strengthen competitiveness in modern agriculture^4^. Previously, a Russian breeding program focusing on common buckwheat with determinate growth habit demonstrated the potential of traditional breeding approaches to improve this crop and developed numerous varieties showing exceptional agronomic performance in Western European contexts^5–8^. However, the yields and economic revenues of these varieties are not yet competitive to major crops, highlighting the urgency of improving common buckwheat using modern genomic techniques^9,10^. To implement state-of-the-art genomic techniques in research and breeding, the availability of high-quality genomic resources, such as reference-level genome assemblies, are required.

Over the past decade, several genome assemblies of common buckwheat have been generated. The first draft genome assembly, with a scaffold N50 of 25 kb, was produced using Illumina short-read sequencing and published in 2016^11^. Five years later, a genome assembly representing the Russian cultivar ‘Dasha’ was reported^12^. This assembly, based on a combination of Illumina paired-end reads and PacBio HiFi reads, showed improved contiguity, with a higher scaffold N50 of 180 kb^12^. In 2023, a chromosome-level genome assembly of a self-compatible, highly homozygous common buckwheat line (PL4) was released^13^. Both Illumina paired-end reads and PacBio HiFi reads were used to assemble PL4, with additional scaffolding using a genetic linkage map, increasing the scaffold N50 to 155 Mb^13^. In the same year, a chromosome-level, haplotype-resolved genome assembly of the Chinese common buckwheat cultivar ‘Xinong9976’ was described^14^. Generated using PacBio HiFi reads and high-throughput chromosome conformation capture (Hi-C) data, this assembly comprised two haplotype sets (hereafter referred to as haplomes), with scaffold N50 values of 156 Mb and 150 Mb^15^, respectively.

Despite these advances, no high-quality, haplotype-resolved genome assembly of a European elite variety is currently available. Besides, using genome assemblies with distinct genetic backgrounds to study European materials may introduce reference bias in downstream analyses and applications. To address this gap and support the improvement of common buckwheat varieties in Europe, we sequenced an F_1_ genotype (named Tuka), derived from the cross between a European elite cultivar, ‘Devyatka’ and a self-compatible variety, ‘Tussi’, following a trio-binning approach^15^. This resulted in high-quality Illumina paired-end sequencing data for both parents, as well as PacBio HiFi and Hi-C data for Tuka. Assembly of these data resulted in a near-complete, chromosome-level, haplotype-resolved assembly comprising two haplomes for Tuka. Furthermore, evidence-based gene prediction using data from short-read RNA sequencing (RNA-seq) and isoform sequencing of full-length transcripts (Iso-seq) of Tuka progeny yielded complete and accurate gene annotations for both haplomes. All sequencing data, assemblies and annotations generated in this study constitute valuable genomic resources that will facilitate downstream analyses and applications, including marker development, genome-wide association studies, genomic selection and gene editing, to advance common buckwheat research and breeding.

## Methods

### Plant materials

A trio of common buckwheat plants was created by crossing one individual of the cultivar ‘Devyatka’ (mother or pollen recipient, high-yielding, self-incompatible with determinate growth habit) with an individual of another variety ‘Tussi’ (father or pollen donor, self-compatible with indeterminate growth habit, Fig. 1). These two original plants are referred to as the parental generation (P), and they were used for Illumina whole-genome short-read sequencing. To cross the two parental plants, a brush was used to transfer pollen from ‘Tussi’ to the style of ‘Devyatka’. The offspring of this cross is referred to as the F_1_ generation. A single F_1_ plant (named Tuka), which is self-compatible with indeterminate growth habit, was selected for whole-genome long-read sequencing and Hi-C sequencing. Furthermore, Tuka was allowed to flower in isolation to ensure self-pollination, generating F_2_ individuals that were subsequently used for RNA-seq and Iso-seq.

**Fig. 1.**
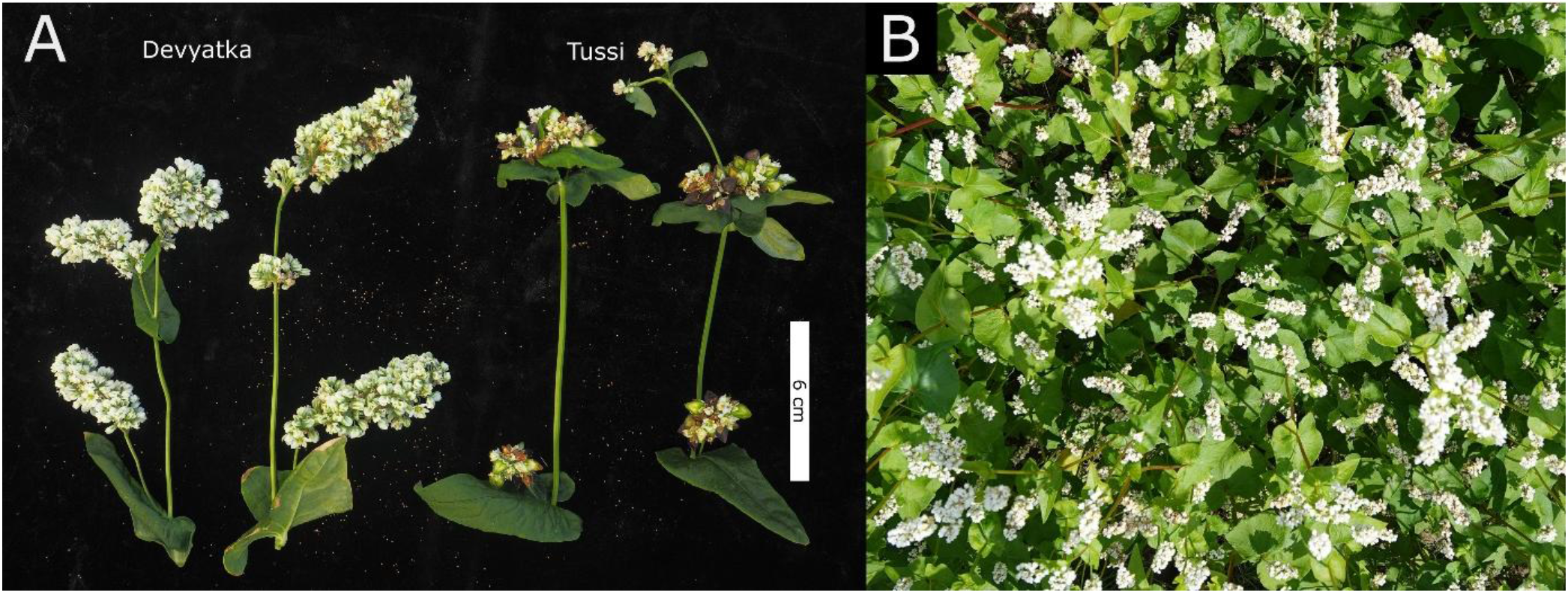
Photographs illustrating the morphology of common buckwheat. (A) Morphology of terminal flower clusters of the cultivar ‘Devyatka’ (left) and the variety ‘Tussi’ (right). (B) Buckwheat hybrids (‘Devyatka’ × ‘Tussi’) in the field. Photo credit: Miguel Angel Loera Sanchez, Sebastian Soyk and Fabian Hess

### Whole-genome short-read sequencing of the parental plants

Young leaves were collected from both parental plants for DNA extraction, and the collected leaves were snap-frozen in liquid nitrogen. DNA was extracted using the DNeasy Plant Mini Kit (Qiagen, Hilden, Germany) following the manufacturer’s instructions. Library construction and sequencing were done at the Functional Genomics Center Zurich (FGCZ). Library construction was performed using the Illumina Truseq Nano protocol, and an Illumina Novaseq 6000 device (Illumina, San Diego, USA) was used for sequencing. In total, 116.58 and 96.76 Gb paired-end reads with a read length of 150 bp were obtained for the maternal and the paternal plant of ‘Devyatka’ and ‘Tussi’, respectively.

### PacBio HiFi sequencing and Hi-C sequencing of Tuka

For PacBio HiFi sequencing, high molecular weight DNA was extracted from young leaves of Tuka using the protocol described by Russo et al.^16^, adapted to omit EDTA from the SDS buffer. Library construction and PacBio HiFi sequencing was done at FGCZ. The HiFi standard protocol for 15-20kb libraries was used to construct two libraries, one of which was sequenced using PacBio Sequel IIe, while the other library was sequenced on the PacBio Revio (PacBio, Menlo Park, CA, USA). In total, 122.83 Gb HiFi reads with a read N50 of 17.94 kb were obtained. The sequencing coverage of the HiFi reads is around 102-fold, assuming a haploid genome size of 1.20 Gb.

The same leaf material as for PacBio HiFi sequencing was used for the construction of a Hi-C library. The Hi-C sample was generated following a previously published protocol^17^ optimized for non-model plant species and using HindIII as restriction enzyme. The library was prepared using a customized protocol based on the KAPA HyperPrep (Roche, Basel, Switzerland) library preparation kit. The Hi-C library was subsequently sequenced at FGCZ using Illumina NovaSeq X Plus (Illumina, San Diego, USA). In total, 236.67 Gb paired-end Hi-C reads with a length of 150 bp were obtained. The sequencing coverage of the Hi-C reads is around 197-fold, assuming a haploid genome size of 1.20 Gb.

### Short-read and long-read transcriptome sequencing

Nineteen F_2_ plants for each of the two different phenotypes (determinate and indeterminate growth habit) were grown for transcriptome sequencing. On the same day, 18 plants (nine per phenotype) were placed in the greenhouse, and 20 plants (ten per phenotype) were placed in the climate chamber. Growing conditions and environmental parameters were standardized across both locations. Each pot contained three liters of a premixed substrate consisting of 25% sterilized soil, 25% bark compost, 20% washed sand and 30% white peat (Ricoter, Aarberg, Switzerland). Plants were exposed to eight hours of light per day, with temperatures maintained between a minimum of 10°C and a maximum of 25°C. At both locations, four tissue types (leaf, flower, root and stem) were collected from all plants of each phenotype. With two technical replicates per tissue type, leaves and flowers were sampled 37 days after sowing, and roots and stems were sampled 56 days after sowing. This resulted in 2 samples per tissue type for each phenotype at each location, yielding 32 samples for subsequent RNA extraction.

Total RNA was extracted from all 32 samples using the NucleoSpin RNA Plus, Mini kit for RNA purification with DNA removal column (MACHEREY-NAGEL, GmbH & Co. KG, Düren, Germany; cat. no. 740984.50). Quality and concentrations of each RNA sample was assessed using a Qubit fluorometer (Thermo-Fisher Scientific, Waltham, MA, USA) and Agilent TapeStation system (Agilent Technologies, Santa Clara, CA, USA) at Genetic Diversity Center (GDC). For RNA-seq, one RNA sample per tissue type for each phenotype and location was selected, yielding 16 samples. The remaining 16 RNA samples were pooled by phenotype to generate two mixed samples comprising all tissue types for Iso-seq. All RNA-seq and Iso-seq libraries were sequenced by Novogene GmbH (Munich, Germany).

For RNA-seq, libraries were prepared using a directional mRNA protocol with poly(A) enrichment and sequenced as paired-end reads (PE150) on an Illumina NovaSeq X Plus platform, generating approximately 6 Gb of raw data per sample. This resulted in 16 RNA-seq datasets, which were merged into a single dataset for downstream gene prediction. In total, 103.44 Gb of RNA-seq data with a read length of 150 bp were generated.

For Iso-seq, the two pooled samples were used to build a single library using the Kinnex protocol (PacBio, Menlo Park, CA, USA). The library was sequenced on a PacBio Revio system to generate HiFi reads, yielding approximately 5 million raw reads per pooled sample. This resulted in two Iso-seq datasets, which were concatenated into a single dataset for downstream gene prediction. In total, 25.20 Gb Iso-seq data were obtained, with a read N50 of 1.67 kb.

### Estimation of genome size and level of heterozygosity

The haploid genome size and the level of heterozygosity of Tuka were estimated using GenomeScope2^18^ v2.0.1 based on k-mers (length of 23 bp), counted from PacBio HiFi reads by using jellyfish^19^ v2.2.10. The estimated haploid genome size was 1.26 Gb, and the estimated level of heterozygosity was 2.06%, suggesting at least two single nucleotide polymorphisms per 100 bp.

### Genome assembly and scaffolding

To generate the haplotype-resolved assembly for Tuka, Hifiasm^20^ v0.19.5-r587 with the trio-binning mode was used based on the following two steps: First, maternal- and paternal-specific k-mers (length of 31 bp) were extracted from the parental Illumina short-read data using yak (yak-0.1 r56, https://github.com/lh3/yak). Second, PacBio HiFi reads of Tuka, together with the maternal- and paternal-specific k-mers, were fed into Hifiasm for assembly, resulting in two contig-level haplomes.

A two-step strategy was applied to scaffold each haplome separately with the following two steps: First, each haplome was anchored to the published chromosome-level genome assembly of common buckwheat, PL4^21^, with RagTag^22^ v2.1.0, resulting in two scaffold-level haplomes. Second, Hi-C reads were mapped to each scaffold-level haplome using Juicer pipeline^23^ v2.0, resulting in two Hi-C contact maps. Then, based on each Hi-C contact map, manual inspection and curation was conducted using JuiceBox^24^ v1.11.08. Finally, with the manually curated Hi-C contact maps, two chromosome-level haplomes, named Tuka_h1 and Tuka_h2, were generated using 3d-DNA^25^ v201008.

### Transposable element annotation

Transposable element (TE) annotation for each haplome was performed using the Extensive de-novo TE Annotator (EDTA^26^ v2.2.2), and 73.96% and 75.16% of Tuka_h1 and Tuka_h2 were identified as TEs, respectively (Table 2).

### Evidence-based protein-coding gene prediction

To avoid over-predicting TEs as protein-coding genes, TEs identified in each haplome were soft-masked by turning bases of TEs lowercase, prior to gene prediction. An evidence-based gene prediction pipeline was used to predict protein-coding genes in each haplome separately with different types of evidence listed below.

#### Evidence 1: transcript alignment

Short reads resulting from RNA-seq were aligned to each haplome using HISAT2^27^ v2.2.1 (parameters: –max-intronlen 20000 –dta -p 16 –rna-strandness RF –fr –no-mixed –no-discordant –omit-sec-seq). PacBio HiFi long reads resulting from Iso-seq were aligned to each haplome using Minimap2^28^ v2.28 (parameters: -x splice:hq -L -2 -G 20000 –secondary=no). Based on the alignments of both RNA-seq and Iso-seq data, transcripts were assembled using StringTie^29^ v3.0.0 with the mixed assembly mode (parameters: –mix -f 0.1 -p 8 -c 3 -j 3 -t -m 100 –rf). Based on the alignment of Iso-seq data, transcripts were also assembled using StringTie v3.0.0 with the long-read-only-assembly mode (parameters: -L -f 0.1 -p 8 -c 3 -j 3 -t -m 100 -s 5 -g 0).

#### Evidence 2: protein alignment

Protein sequences from the UniRef90 Embryophyta protein database were aligned to each haplome using Miniprot^30^ v0.16 (parameters: -G 20000 --gff-only --outc 0.8). Proteins with both alignment coverage and identity >= 80% were retained for further gene prediction.

#### Evidence 3: *ab initio* gene prediction

BRAKER3^31^ v3.0.8 was used for de novo gene prediction based on the RNA-seq read alignment obtained when generating evidence 1. Galba^32^ v1.0.11 was used for de novo gene prediction with the proteins predicted by TD2 v5.7.1, which predicted proteins based on the mixed transcript assembly obtained when generating evidence 1.

#### Evidence 4: other prediction

The proteins predicted by TD2^33^, when generating evidence 3, were used as additional protein evidence. UniRef90 Embryophyta was used as a reference protein database with TD2 to filter non-coding proteins.

All types of evidence, i.e., transcript alignment, protein alignment, *ab initio* gene prediction and other prediction, were provided to EVidenceModeler (EVM^34^, v2.1.0) to build consensus gene models. This resulted in 38,779 and 35,884 gene models for Tuka_h1 and Tuka_h2, respectively (Table 1).

**Table 1.**
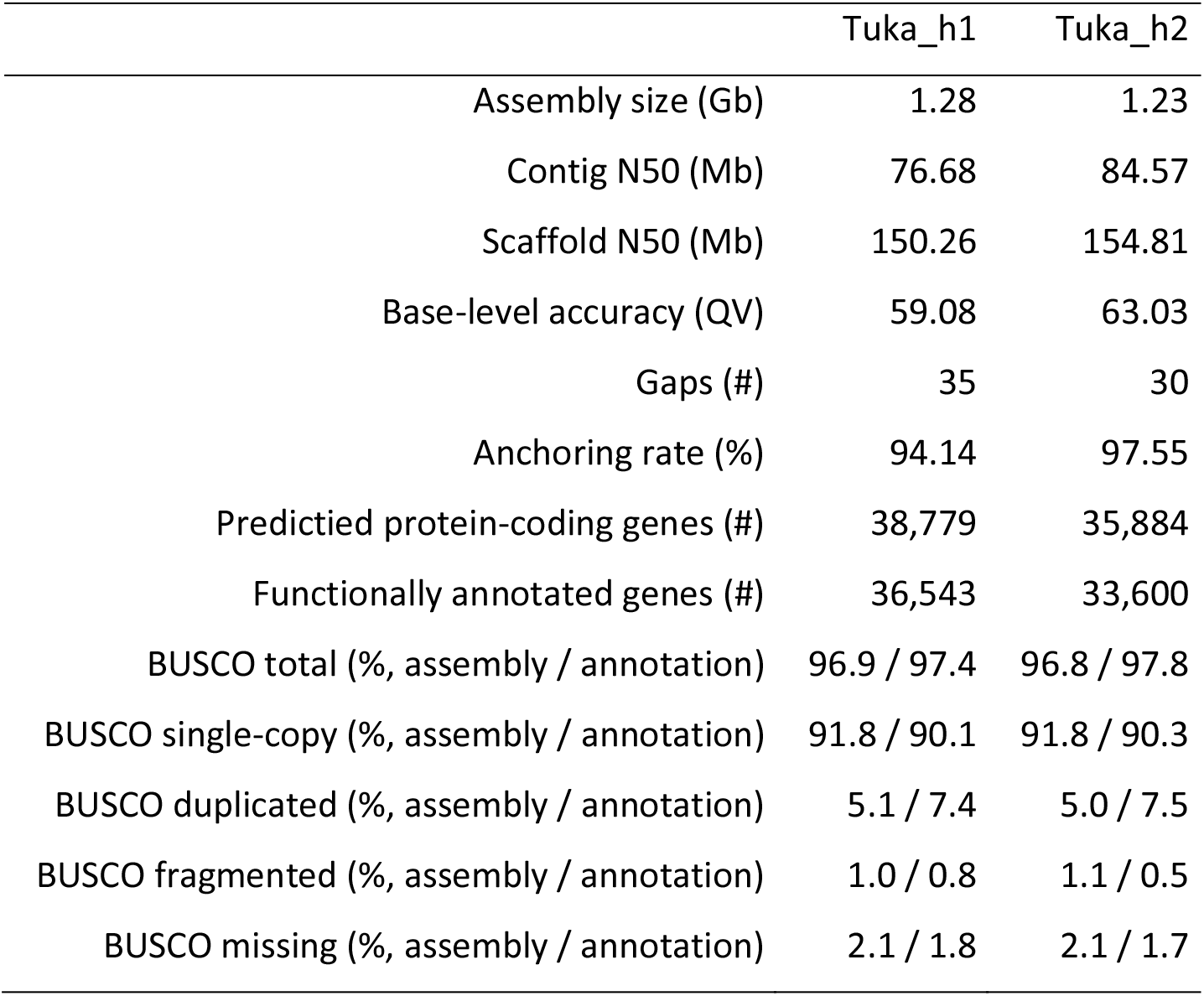
Genome assembly and annotation statistics.

### Functional gene annotation

To add a human-readable description of the gene function to each gene model, following EviAnn^35^ v2.0.1, gene functions of proteins in public database were transferred to the gene models using a custom script conducting the following steps: First, the predicted protein-coding genes (hereafter referred to as query proteins) were aligned against the UniRef100 (Embryophyta) database (hereafter referred to as reference proteins) using MMseqs2^36^ v17.b804f. Second, if a reciprocal best hit (RBH) were found for a query protein, gene function of the reference protein was assigned to the query protein. If no RBH was found, gene function from the reference protein of the best BLAST hit (BBH) was assigned to the query protein. Finally, if neither RBH nor BBH were detected, no function was transferred to the query protein. The description of gene function was added as a tag into the ninth column of the annotation file (gff file). In total, 36,543 and 33,600 genes were functionally annotated in Tuka_h1 and Tuka_h2, respectively (Table 1).

### Quality control of genome assembly and annotation

Basic assembly statistics, including assembly size, contig N50 and scaffold N50, were calculated using SeqKit2^37^ v2.9.0 (parameters: stats -a -b). The presence of telomeres was detected using tidk^38^ v0.2.65 by searching for “GAAACC”, as suggested by Li et al^39^. The anchoring rate was calculated by dividing the size of pseudo-chromosomes of each haplome to the total size of each haplome. Gaps in each haplome were identified by searching for contiguous sequences of Ns using SeqKit2 v2.9.0 (parameters: locate -p N+ -G -P -r -i –bed). To check the correctness of the assembly, k-mers (length of 23 bp) from each haplome were compared with k-mers from the total PacBio HiFi reads using KAT^40^ v2.4.2. This resulted in the number of total k-mers of each haplome and the number of shared k-mers between each haplome and the reads, which were used to estimate the base-level accuracy, following the formula of consensus quality estimation described in Merqury^41^. To check whether the two haplotypes have been properly separated, PacBio HiFi reads were aligned to the diploid assembly (the concatenation of Tuka_h1 and Tuka_h2) using Minimap2 v2.26-r1175, and based on the read alignment, median alignment coverage was calculated for each 1 Mb window using Mosdepth^42^ v0.3.3. To check the completeness of the gene space of both haplomes, BUSCO^43^ v5.3.2 with 1,614 BUSCO genes in embryophyta_odb10 database was used. To check the synteny and orientation between two homologous pseudo-chromosomes, whole-genome alignment was conducted between Tuka_h1 and Tuka_h2 using FastGA^44^ v1.3.1.

To check the completeness of the gene prediction, BUSCO v5.8.2 with 2,026 BUSCO genes in embryophyta_odb12 database was used. To visualize the distribution of genes and TEs in the pseudo-chromosomes of both haplomes, the proportion of bases belonging to genes or TEs in every 1 Mb window was calculated using samtools^45^ v1.3.1 and bedtools^46^ v2.31.1 with the following steps: First, sizes of pseudo-chromosomes were calculated using samtools faidx, resulting in the .fai index file. Then, the first two columns in the .fai index file were extracted and stored in a .genome file required by bedtools. Next, a .bed file with 1 Mb windows of all pseudo-chromosomes was generated using bedtools makewindows. Finally, the proportion of bases belonging to genes or TEs in every 1 Mb window was calculated using bedtools coverage based on the .bed window file, the gene annotation and TE annotation files. Following the same steps, the proportion of non-repetitive sequences in every 1 Mb window was also visualized, and the non-repetitive regions in each haplome were detected by using KAT v2.4.2 (parameters: kat sect -E -F -M 2) with the same haplome as both reference and query. Following Xu et al.^47^, the 1 Mb window showing the lowest proportion of non-repetitive sequences in the middle of each pseudo-chromosome together with 1 Mb window upstream and downstream was defined as the putative centromeric region. Genomic features, including the length of pseudo-chromosomes, position of gaps, telomeres and centromeres, proportion of non-repetitive sequences, genes and TEs, PacBio HiFi read alignment depth and the whole-genome alignment between Tuka_h1 and Tuka_h2, were visualized with a circos plot using circlize^48^ v0.4.16.

### Data Records

Illumina whole-genome sequencing data of both the maternal (from the cultivar ‘Devyatka’, tag ID: FE213) and the paternal (from the variety ‘Tussi’, tag ID: FE228) plant have been deposited in the European Nucleotide Archive (ENA) at EMBL-EBI under accession number PRJEB103817^49^. PacBio HiFi, Hi-C, RNA-seq and Iso-seq data of Tuka have been deposited in ENA at EMBL-EBI under accession number PRJEB103817^49^. The haplomes of Tuka, Tuka_h1^50^ and Tuka_h2^51^, and the corresponding TE and gene annotation files, have been deposited in ENA at EMBL-EBI under accession number PRJEB103817^49^.

### Technical Validation

The two haplomes, Tuka_h1 and Tuka_h2, showed a total size of 1.28 and 1.23 Gb, respectively (Table 1), and ∼1.20 Gb sequences were anchored to eight pseudo-chromosomes in each haplome, corresponding to a high anchoring rate of 94.14% and 97.55%, respectively (Table 1).

High contiguity was observed for Tuka_h1 and Tuka_h2, as indicated by the high contig N50 of 76.68 and 84.57 Mb, respectively, and the chromosome-level scaffold N50 of 150.26 and 154.81 Mb, respectively (Table 1). Only 35 and 30 gaps were detected in Tuka_h1 and Tuka_h2, respectively (Table 1, Fig. 2), and no gaps were found in chr3 of Tuka_h1 (Fig. 2). Telomeres were found at both ends of all pseudo-chromosomes, except chr5, chr7 and chr8 of both haplomes, where signals of telomeres were only observed at one end (Fig. 2, Table 3).

**Table 2.**
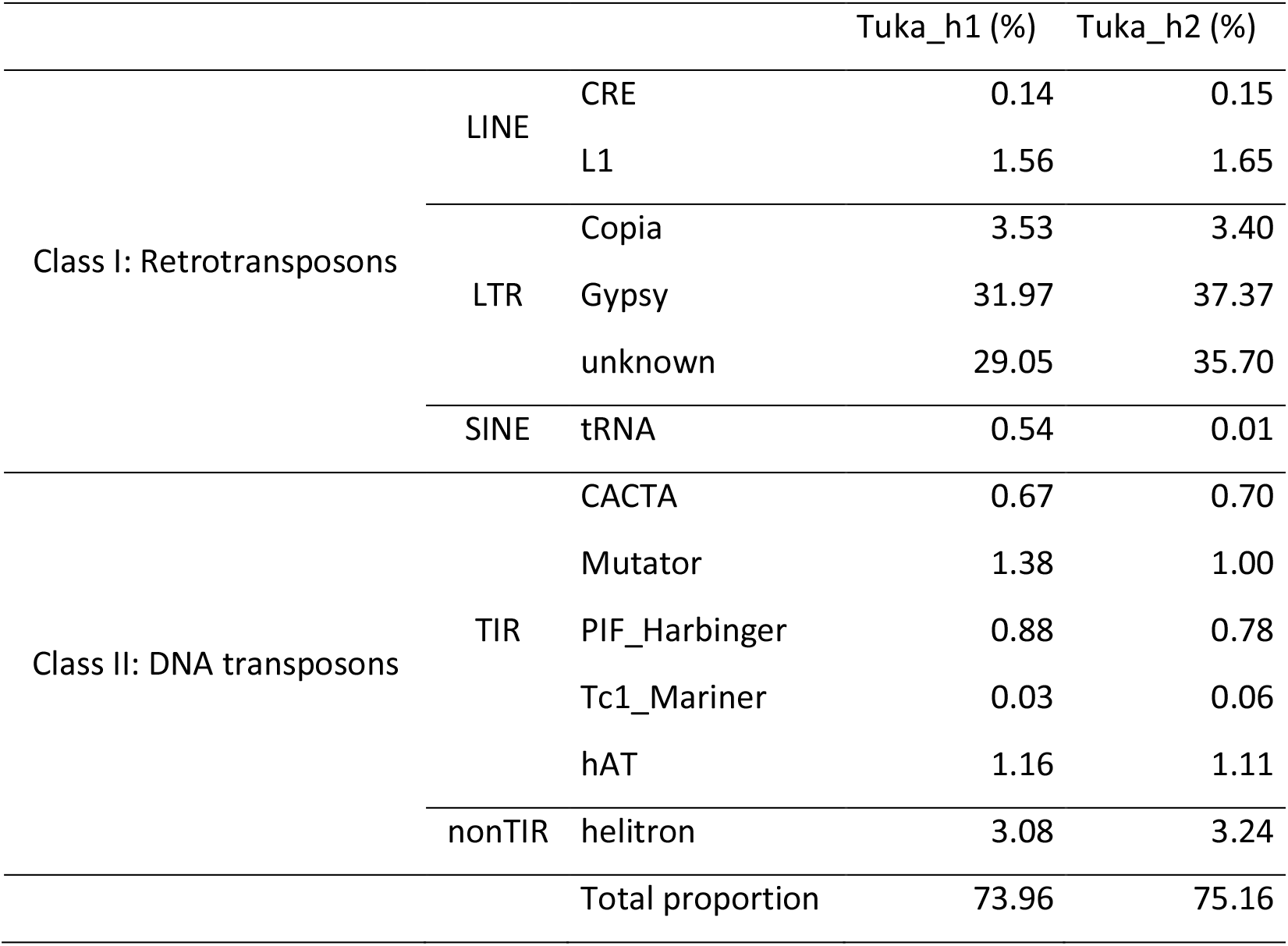
Proportion of transposable elements in each haplome.

**Table 3.** Estimated positions of centromeres and telomeres.

| Chromosome | Centromere<br>start (bp) | Centromere<br>end (bp) | Left telomere<br>start (bp) | Left telomere<br>end (bp) | Right telomere<br>start (bp) | Right telomere<br>end (bp) |
| --- | --- | --- | --- | --- | --- | --- |
| chr1_h1 | 75,000,000 | 78,000,000 | 0 | 10,000 | 162,260,000 | 162,267,147 |
| chr2_h1 | 87,000,000 | 90,000,000 | 0 | 10,000 | 161,200,000 | 161,215,231 |
| chr3_h1 | 75,000,000 | 78,000,000 | 0 | 10,000 | 150,240,000 | 150,258,585 |
| chr4_h1 | 86,000,000 | 89,000,000 | 0 | 20,000 | 156,510,000 | 156,516,947 |
| chr5_h1 | 69,000,000 | 72,000,000 | - | - | 153,410,000 | 153,430,000 |
| chr6_h1 | 80,000,000 | 83,000,000 | 0 | 20,000 | 145,180,000 | 145,200,165 |
| chr7_h1 | 67,000,000 | 70,000,000 | 0 | 10,000 | - | - |
| chr8_h1 | 58,000,000 | 61,000,000 | - | - | 136,510,000 | 136,521,069 |
| chr1_h2 | 77,000,000 | 80,000,000 | 0 | 10,000 | 163,630,000 | 163,642,019 |
| chr2_h2 | 88,000,000 | 91,000,000 | 0 | 10,000 | 160,390,000 | 160,396,531 |
| chr3_h2 | 78,000,000 | 81,000,000 | 80,000 | 90,000 | 154,800,000 | 154,806,493 |
| chr4_h2 | 84,000,000 | 87,000,000 | 0 | 10,000 | 155,970,000 | 155,988,789 |
| chr5_h2 | 68,000,000 | 71,000,000 | - | - | 151,110,000 | 151,119,348 |
| chr6_h2 | 78,000,000 | 81,000,000 | 0 | 10,000 | 141,400,000 | 141,409,865 |
| chr7_h2 | 65,000,000 | 68,000,000 | 0 | 10,000 | - | - |
| chr8_h2 | 58,000,000 | 61,000,000 | - | - | 133,930,000 | 133,941,421 |

**Fig. 2.**
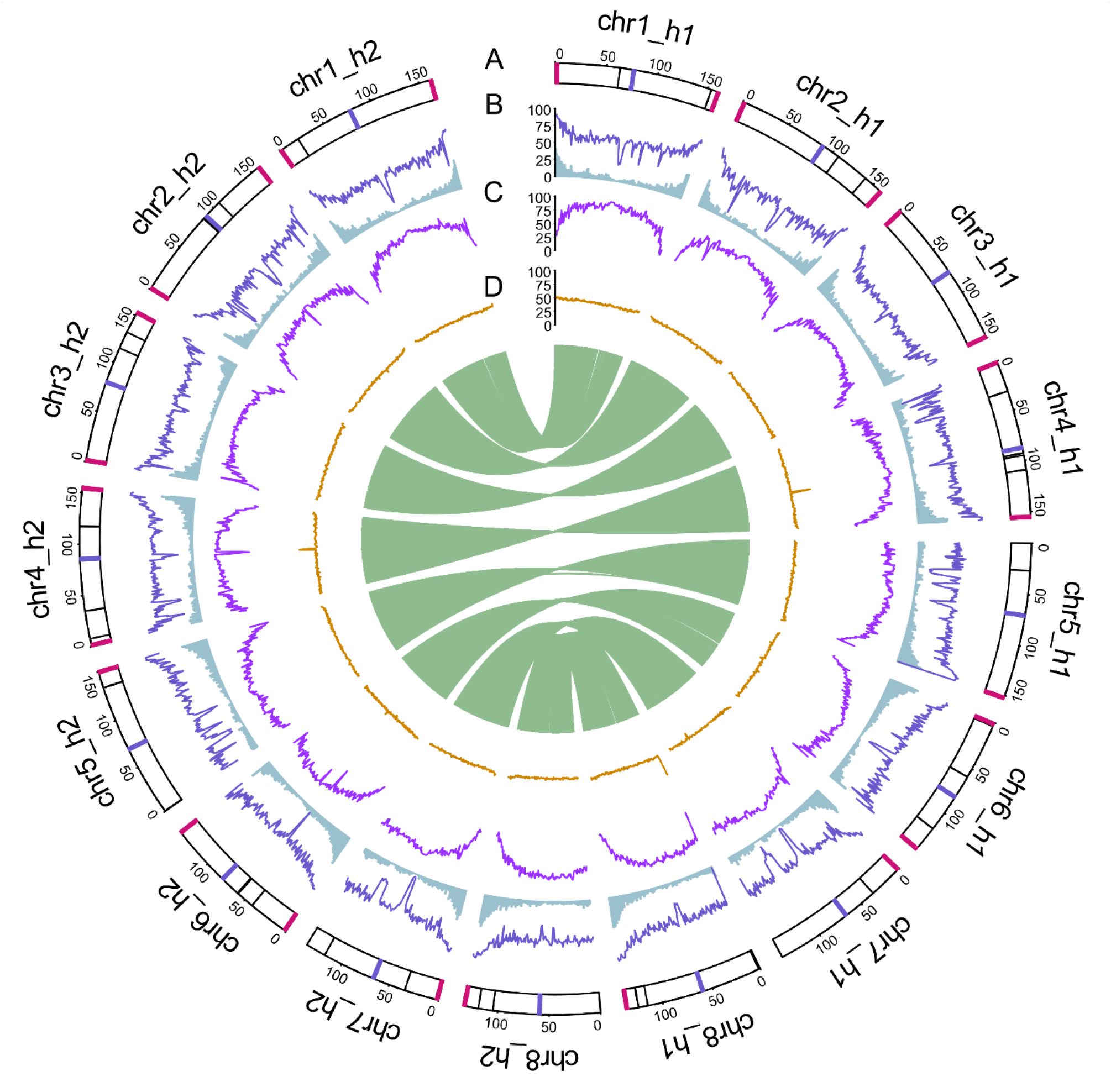
Circos plot showing the genomic features of Tuka. (A) Pseudo-chromosomes (Mb) of the two haplomes, Tuka_h1 and Tuka_h2. Red indicates telomeres. Blue indicates putative centromeres. Black lines indicate gaps. (B) Dark blue and light blue indicate the proportion of non-repetitive sequences and genes in every 1 Mb window. (C) Proportion of transposable elements in every 1 Mb window. (D) PacBio HiFi read alignment depth. Green links in the center indicate whole-genome alignment between Tuka_h1 and Tuka_h2. Only alignments between two homologous pseudo-chromosomes with an alignment length over 10 kb were visualized.

Each haplome was found to be highly complete, with a total complete BUSCO score of 96.9 and 96.8 for Tuka_h1 and Tuka_h2, respectively (Table 1). Proteins predicted from each haplome also showed a high completeness, with a total complete BUSCO score of 97.4 and 97.8 for Tuka_h1 and Tuka_h2, respectively (Table 1).

Each haplome was found to be correctly representing a haploid genome, as revealed by the k-mer comparison analyses with KAT v2.4.2 (Fig. 3), and each haplome showed correct structure for all pseudo-chromosomes, as indicated by its Hi-C contact map (Fig. 4). The whole-genome alignment between Tuka_h1 and Tuka_h2 showed a high synteny and a consistent orientation between pairs of homologous pseudo-chromosomes (Fig. 2). A high base-level accuracy was observed for each haplome, with a high quality value (QV) of 59.08 and 63.03 for Tuka_h1 and Tuka_h2, respectively (Table 1), suggesting one putative sequencing error in every 1 Mb.

**Fig. 3.**
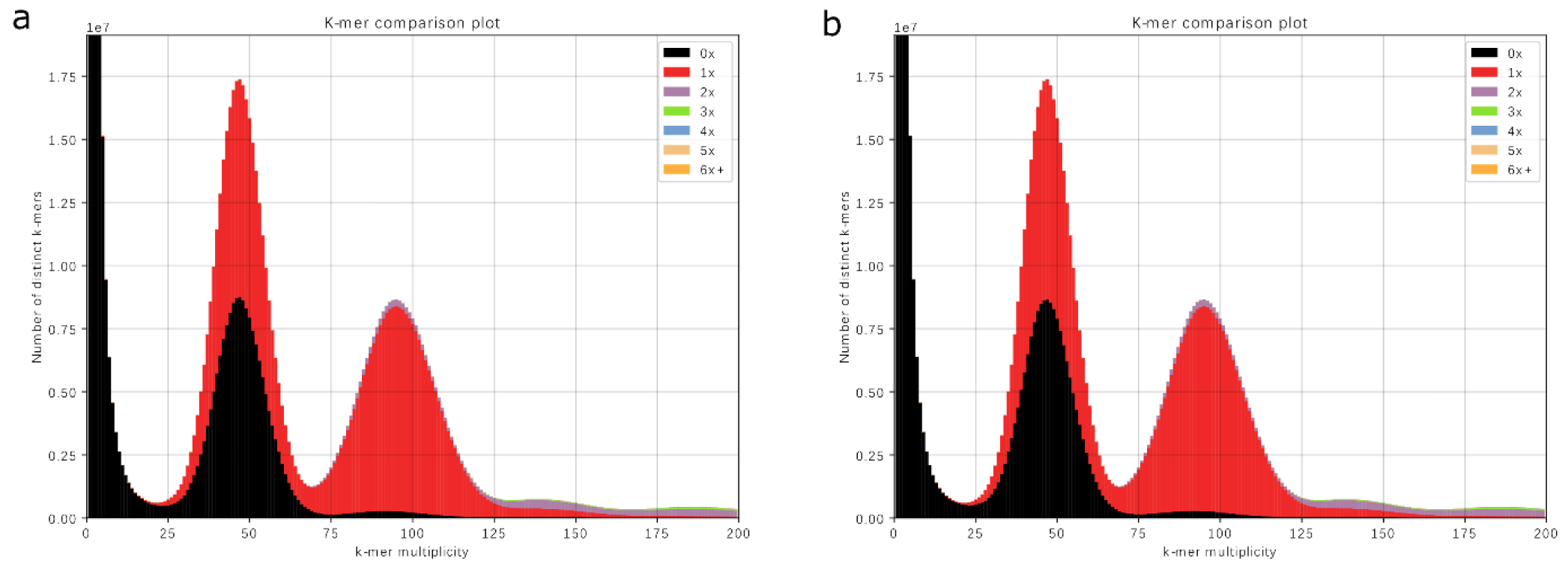
K-mer comparison between each haplome and PacBio HiFi data. (a) K-mer comparison between Tuka_h1 and PacBio HiFi reads. (b) K-mer comparison between Tuka_h2 and PacBio HiFi reads. In each plot, two peaks were observed, with the first and second peak indicating k-mers from heterozygous and homozygous regions, respectively. For a correct haploid assembly, the first peak is expected to show half black and half red, meaning that only k-mers of one haplotype at heterozygous regions are present in the assembly and are only present once. The second peak is expected to be fully red, meaning that k-mers from homozygous regions are all present in the assembly and are only present once.

**Fig. 4.**
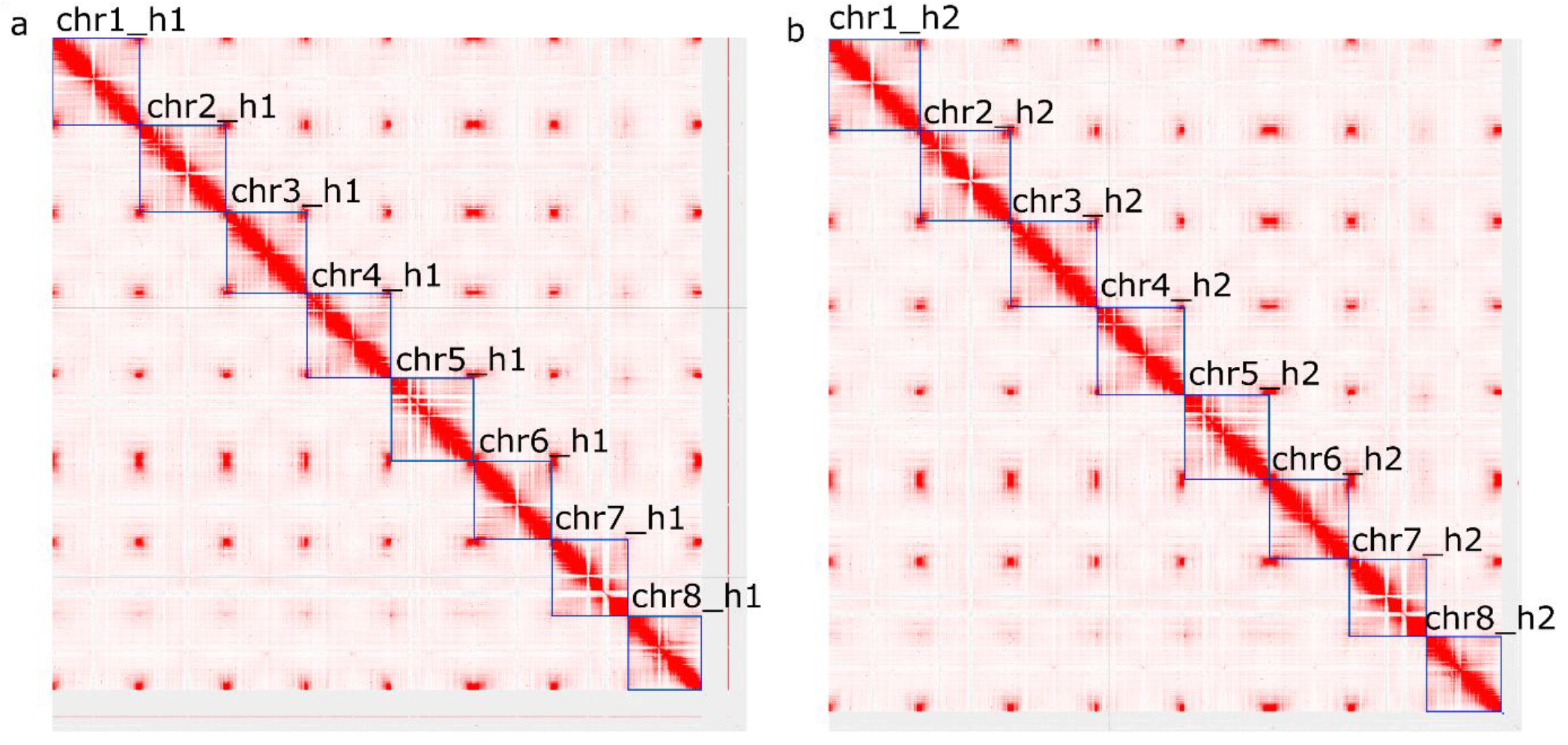
High-throughput chromosome conformation capture (Hi-C) contact map of Tuka_h1 (a) and Tuka_h2 (b). In both Hi-C contact maps, each blue box represents one pseudo-chromosome, with the corresponding name shown above the box.

Each haplome showed a consistent ∼50-fold PacBio HiFi read alignment coverage for all pseudo-chromosomes (Fig. 2), when mapping a total of ∼102-fold PacBio HiFi reads to the diploid assembly. This suggests that haplotypes were properly separated or phased during assembly. Notably, a few genomic regions showed a much higher PacBio HiFi read alignment coverage, such as chr4_h1, chr8_h1 and chr5_h2, where a high peak of coverage was observed (Fig. 2). This suggests that these regions might be highly repetitive and were likely collapsed during assembly.

Genes were found to be not evenly distributed in pseudo-chromosomes. A higher proportion (up to 48.74%) of genes was observed at the ends of the pseudo-chromosomes, when compared to the centromeric regions (Fig. 2, Table 3). TEs showed the highest proportion (up to 99.59%) in the centromeric regions of pseudo-chromosomes, and the proportion of TEs drops gradually when moving from the centromeric region towards the telomeric region (Fig. 2).

## Code Availability

Code for assembly, scaffolding and technical validation is available at https://github.com/Yutang-ETH/Buckwheat_Tuka.

## Author Contributions

B.S., R.K., F.H. conceived this study, and B.S. acquired funding. F.H. prepared and collected all plant materials, conducted whole-genome sequencing and wrote the manuscript. Y.T.C. performed genome assembly, scaffolding and assembly quality check analyses and wrote, together with F.H., the manuscript. M.E.L. extracted RNA, conducted RNA-seq and Iso-seq and contributed to manuscript writing. A.C. performed TE annotation, gene prediction and quality check analyses for the predicted genes. I.S.S. extracted DNA for whole-genome sequencing. S.G. constructed the Hi-C library. B.S. and R.K. supervised this work and edited the manuscript. All authors reviewed and approved the manuscript for publication.

## Acknowledgements

We sincerely thank Verena Knorst from the Molecular Plant Breeding group at ETH Zurich for carefully maintaining the plant materials. We gratefully acknowledge FGCZ for providing their sequencing services and GDC at ETH Zurich for providing their facilities for DNA and RNA quality check. We thank Dr. Michelle Nay and Dr. Lukas Kronenberg for their input and inspiration during the conceptualization of this study and appreciate Dr. Steven Yates for his helpful and constructive suggestions on this manuscript. We sincerely thank ISG-HEST at ETH Zurich for providing computational resources as well as their IT service for this work.

